# *In silico* modelling and characterization of Epstein–Barr virus LMP1 protein

**DOI:** 10.1101/2024.09.30.615964

**Authors:** Dayang-Sharyati D.A. Salam, Kavinda Kashi Juliyan Gunasinghe, Hwang Siaw San, Irine Runnie Henry Ginjom, Xavier Chee Wezen, Taufiq Rahman

## Abstract

Latent membrane protein 1 (LMP1) plays a crucial role in Epstein-Barr virus (EBV)’s ability to establish latency and is involved in the development and progression of EBV-associated cancers. Additionally, EBV-infected cells affect the immune responses, making it challenging for immune system to eliminate. Due to the aforementioned reasons, it is important to understand the structural features of LMP1 which is essential for the development of novel cancer therapies that target its signaling pathways. To date, there is no complete LMP1 protein structure therefore in our work, we modeled the full-length LMP1 containing the short cytoplasmic N-terminus, six transmembrane domains and a long-simulated C-terminus. Our model showed good stability and protein compactness evaluated through accelerated-Molecular Dynamics where the conformational ensemble exhibited compact folds, particularly in the transmembrane domains. Our results suggest that specific domains or motifs, predominantly in the C-terminus domain of LMP1 show promise as potential drug targets. As a whole, our work provides insights on key strucutral features of LMP1 that will allow the development of novel LMP1 therapies.

## 1.1 Introduction

Epstein-Barr Virus (EBV) is a double-stranded DNA Virus that is commonly contracted by humans. It is estimated that over 90% of the world population had contracted EBV at some point in their lives.^1^ Latent membrane protein 1 (LMP1) is a protein encoded by the EBV and plays a crucial role in EBV’s ability to establish latency. In this state, the virus persists in infected cells without actively replicating. It is a viral oncogene that promotes cell proliferation, survival, and migration while inhibiting apoptosis.^2^ LMP1 plays a critical role in developing and progressing EBV-associated cancers. EBV can cause a range of diseases that include infectious mononucleosis, Burkitt lymphoma, nasopharyngeal carcinoma, and Hodgkin lymphoma.^3^ It can also alter the immune response to EBV-infected cells, making it more difficult for the immune system to eliminate. This is due to EBV latency, immune evasion, and cellular transformation. When EBV enters a latent phase, the virus is not actively replicating, but its DNA remains in the cell. This makes it harder for the immune system to detect and destroy the infected cell.^4-5^ EBV then encodes proteins interfering with the immune system’s ability to recognize and kill infected cells. In some cases, EBV can transform a B cell into a cancerous cell. It is a multifunctional protein that can mimic the signalling of several cellular proteins, including CD40, B-cell receptor, and tumour necrosis factor receptor.^6-8^ This allows LMP1 to activate various cellular signalling pathways, including NF-κB and MAPK.^9-11^ Therefore, targeting LMP1 with methods such as monoclonal antibodies, small molecule inhibitors, and gene therapy presents a promising strategy for treating EBV-related pathogenesis.

The structure of LMP1 has provided essential insights into how LMP1 mediates its oncogenic and transforming activities. LMP1 refers to the EBV latent membrane protein type I with a short cytoplasmic N-terminus (NTER), six transmembrane domains (TMD), and a long cytoplasmic C-terminus.^12^ The TMD of LMP1 forms a hexagonal barrel structure, similar to the design of other type I transmembrane proteins. TMD anchors the protein to the cell membrane, allowing it to interact with other membrane-associated proteins.^13^ The NTER domain is short (24 amino acids), located on the cytoplasmic side of the membrane, and contains binding sites for various cellular proteins. Meanwhile, the C-terminus contains cytoplasmic activation regions (CTAR), which mediate the oncogenic and transforming activities of LMP1.^14^ The domain is large (200 amino acids) and located on the cytoplasmic side of the membrane containing several functional motifs involved in signal transduction.

The cytoplasmic NTER and CTAR of LMP1 contain disordered regions important for LMP1’s ability to interact with multiple cellular proteins and activate various signalling pathways.^15^ These regions lack a fixed, stable structure and may become more structured upon interaction with other molecules. The extent of intrinsic disorder varies across different LMP1 domains and can differ depending on the specific prediction method used. A study suggested that LMP1 is an intrinsic disordered protein, and our previous study also showed that up to 40% of the LMP1 protein might be intrinsically disordered.^16-17^ To date, there is no crystal structure of LMP1 available; as such, we attempted to model structure of LMP1 to guide future drug design against LMP1 using *in silico* methods.

In silico methods, such as protein prediction using AlphaFold2, has become a key aspect in understanding protein structure and dynamics. AlphaFold2 is a protein structure prediction tool developed by DeepMind.^18^ It uses a deep learning method to predict the 3D structure of a protein from its amino acid sequence. This breakthrough technology has several critical applications. Firstly, it aids in understanding protein function at a molecular level, which is crucial in various biological studies.^19^ Additionally, AlphaFold2 expedites the drug discovery process by providing valuable insights into the structure of proteins targeted by drugs.^20^ Lastly, the ability to predict protein structures accurately.^21^

Additionally, Molecular Dynamics (MD) simulations are a valuable computational technique for studying protein behaviour at the atomic level. This method uses classical mechanics to simulate protein dynamics by treating atoms as tiny balls and calculating their interactions based on their positions and forces. MD simulations require powerful computers or specialised hardware due to their computational complexity. They provide insights into protein folding, ligand binding, protein-protein interactions, and protein stability.^22^ While AlphaFold2 predicts protein structures, MD simulations reveal how these structures change and function. In our work, we used AlphaFold2 and MD simulations to understand the protein dynamics of LMP1. This paper is the first to present the complete predicted structure of LMP1 protein MD simulation.

## 2. Results & Discussions

### 2.1. Evaluation of the LMP1 protein structure

First, we used protein structure prediction tools namely AlphaFold2 and GalaxyWeb to refine our LMP1 protein model that we constructed previously.^16^ We selected the top model based on the models that showed high pLDDT values on the AlphaFold2 and Galaxy Web. Based on our model, we observed that the residues between 35-200 showed high accuracy where some models (Model 3 and Model 6) showed a pLDDT value exceeding 90 (Figure 1). These residue ranges are in the TMD of LMP1. A study by Veit et al. produced similar results when they predicted the structure of Porcine Respiratory and Reproductive Syndrome Virus dimer using AlphaFold2.^23^ However, the N-terminal domain (residue from number 0-35) and the CTAR (residue of more than 200) have values lower than 50 suggesting that the prediction on these ambiguous regions is not experimentally conclusive since there is not much structural data available.

**Figure 1.**
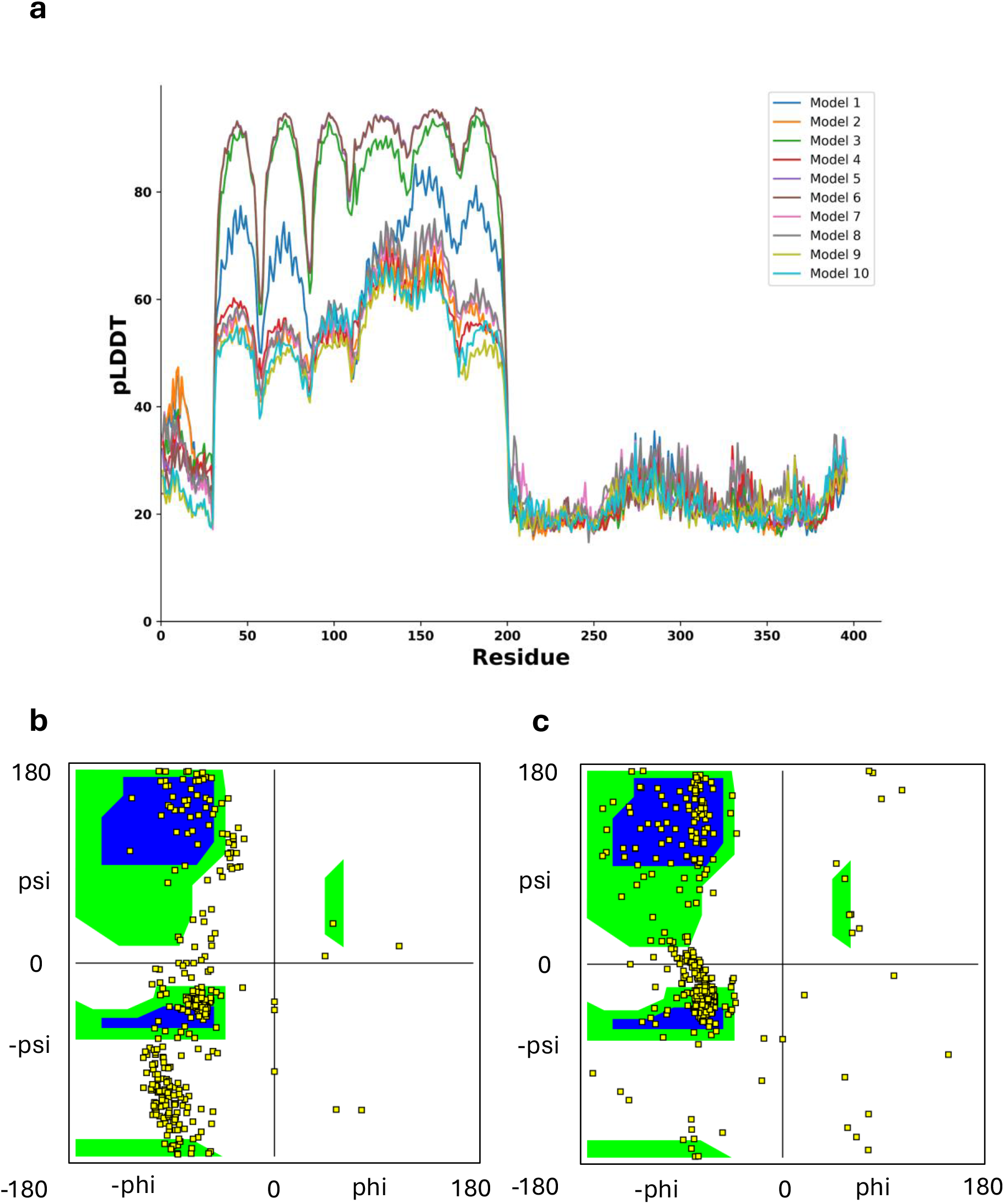
**(a)** Predicted local distance difference test (pLDDT) score per position for the ten predicted models generated by Alphafold2. **(b)** Ramachandran plot for predicted protein Model 3 of Alphafold2. **(c)** Ramachandran plot for our predicted LMP1 protein structure. The Ramachandran Plot showing two core regions (blue colour) and three allowed regions in the three separate boxes (green colour). The beta-sheet region occupies the left-top box, the alpha-helix at the lower-left box and the left-handed helix at the top-right box.

We also used the confidence score of the prediction (between 0 and 1) to assess LMP-1 model quality. The confidence scores indicates how confident AlphaFold2 is in its prediction involving different aspects of protein structure and function, particularly Inferred Post-translational Modifications (IPTM) and Post-translational Modifications (PTM). IPTM are modifications to a protein that occur after it has been synthesised. In the context of AlphaFold2 or any protein structure prediction tool, IPTM would refer to the predicted or inferred post-translational modifications based on the protein’s sequence and other factors. Meanwhile, PTM are actual modifications that occur to proteins after they are translated from mRNA. PTMs regulate protein function, stability, localisation, and interactions with other molecules. They can significantly impact a protein’s structure and activity. AlphaFold2 specifically, it predicts the 3D structure of proteins based on their amino acid sequences, considering evolutionary information and other data. While AlphaFold2 can predict the 3D structure of proteins with high accuracy, it does not explicitly predict post-translational modifications. All the ten model multimers predicted by AlphaFold2 ranged from 0.108 to 0.152 suggesting that the predicted model is naturally disordered or lack sufficient information. Based on the ranking, model 3 has the best score with 0.152 confidence. The graph suggests that the high PLDDT coupled indicate the better-predicted model performance of LMP1 model 3, similar to our LMP1 predicted structure (Figure 1a). A Ramachandran plot was also plotted to showcase both resemblances through the Phi (Φ) and Psi (Ψ) angles, which define a protein’s backbone geometry. Our LMP1 model showed that most of the residues fall between the core regions of the beta-sheet, alpha-helix and left-handed helix regions representing the most energetically favoured conformations (Figure 1b-c). These conformations consisted mainly of the left-handed alpha-helix and beta-sheet structures as exhibit in the TMD of LMP1.

Previous studies on LMP1 suggested that the dimeric form of protein structure is associated with its raft localization and activation and that it is active only in its oligomeric form, specifically the dimeric and trimeric forms.^24-25^ The oligomeric form of LMP1 is critical for its function and activation, and its structure and interactions are essential for LMP1 role in various cellular processes. Therefore, we also investigated LMP1 protein structure using a program available on GalaxyWeb called GalaxyHomomer. The software is used to predict the structure of proteins composed of identical subunits known as homo-oligomers. These proteins are generated when individual protein chains, monomers, come together. GalaxyHomomer includes extra processes to improve the accuracy of the projected structure. This includes modelling the protein’s flexible sections and revising its overall structure. The ab initio docking results of GalaxyHomomer suggested that our LMP1 protein structure consisted of dimeric and trimeric structure conformation such as homo-oligomer (Supplementary Table 1). With the highest docking score value of 2,350.543 and interface area of 2,279.7 (in Å^2^), the predicted model 1 structure consisting of dimer units was similar to the model indicated through AlphaFold2 (Supplementary Figure 1a-b). After the validation of our predicted LMP1 protein structure, we embedded the LMP1 protein in membrane as described in the Computational Methods section. There are various examples of validation in computational biomechanics, emphasizing the importance of experimental data in validating models. We validated our models with the intention of applying predictions to further analysis and experimentation, particularly in the context of patient outcomes. However, comparing model predictions with experimental results to establish credibility in computational modelling is also important.

### 2.2. Simulating LMP1 using accelerated MD simulation

After evaluating the predicted LMP1 protein structure, we simulated the protein using accelerated MD (aMD) simulation for 500 ns in triplicates. The aMD is designed to accelerate the sampling of the phase space by reducing energy barriers, making it more efficient for studying complex systems like protein folding. The force field we used for the aMD simulation was the ff99SBdisp/tip4pd forcefield, originally developed for folded and disordered protein by Robustelli P. *et al*.^26^ There was a recent study discussing the use of ff19SB/OPC forcefield for disordered protein aMD simulation, however, the study suggested the forcefield for the intrinsically disordered protein of less than 50 amino acids.^27^ In our study, our LMP1 protein structure is more than 50 amino acids; hence, we used the ff99SBdisp/tip4pd field.

The aMD simulation revealed several noteworthy observations among LMP1 protein domains or regions. The NTER domain showed root mean squared deviation (RMSD) and radius of gyration (Rg) reached a plateau at around the values of 10 Å indicating a stable structure (Figure 2a-c). NTER experience structural fluctuations at residue Pro10 and Pro20 causing spike in the root mean square fluctuation (RMSF) graph (Figure 2b). The consistent radius of gyration (Rg) value exhibit stability and folded structure of the protein (Figure 2c). The NTER domain’s location may experience these fluctuations, suggesting functional flexibility, such as protein-ligand binding interfaces.

**Figure 2.**
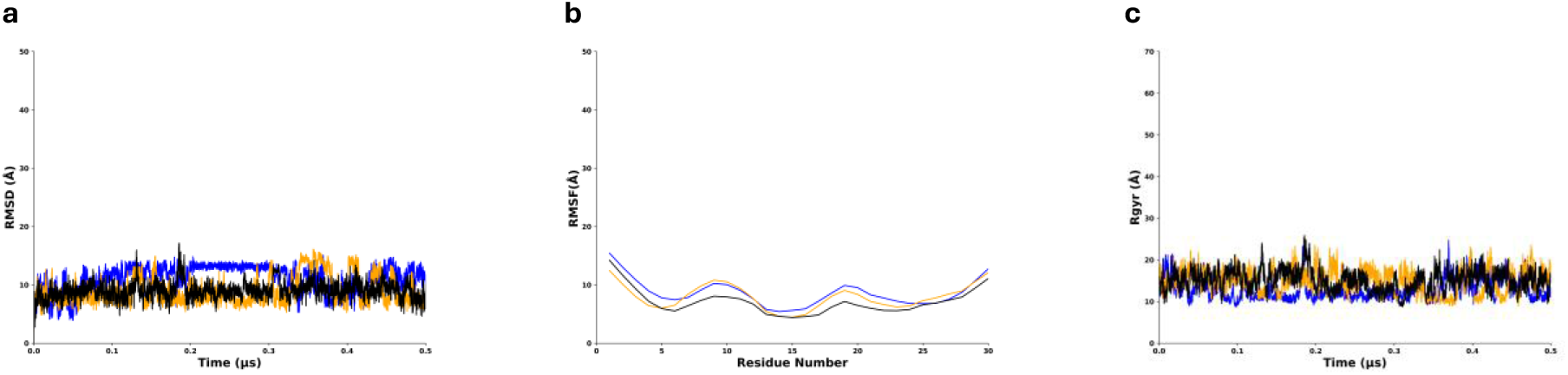
Behaviour of N-terminal regions of LMP1 in MD simulations showing values around 25 Å or lesser for all the three graphs. Replicate 1 is in blue, replicate 2 is in orange, replicate 3 is in black. **(a)** Root mean square deviation (RMSD) graph. **(b)** Root mean square fluctuation (RMSF) graph. **(c)** Radius of gyration (Rgyr) plot.

We noted that the TMD from replicate 1 has a lower threshold from replicates 2 and 3 which have similar values in RMSD, RMSF and Rg. The TMD reached a plateau with an average RMSD value of 15 Å and remains low and steady throughout the simulation indicating convergence to a structure (Figure 3a). The TMD RMSF is 15 Å with multiple fluctuations at residue Gly30, Phe50, Leu80, Leu120 and Leu140, indicating structural variations or flexibility in certain areas, such as flexible loops, protein termini, or solvent-exposed areas (Figure 3b). The Rg value is around 6 Å and consistent throughout, indicating a more folded protein (Figure 3c).

**Figure 3.**
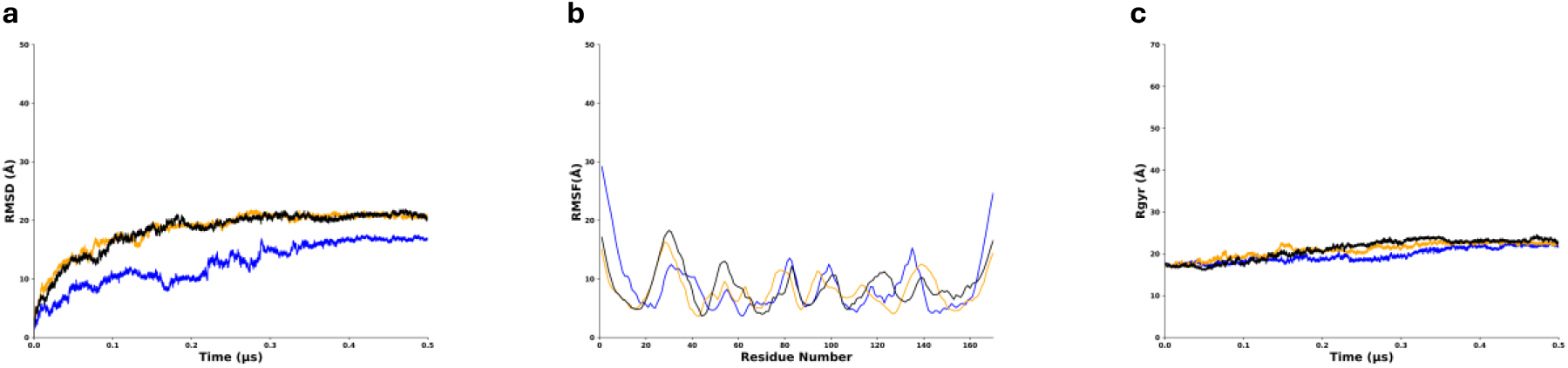
Behaviour of Transmembrane domain of LMP1 in MD simulations showing Replicate 1 RMSD has a lower threshold compared to replicates 2 and 3. Replicate 1 is in blue, replicate 2 is in orange, replicate 3 is in black. **(a)** Root mean square deviation (RMSD) graph. **(b)** Root mean square fluctuation (RMSF) graph. **(c)** Radius of gyration (Rgyr) plot.

In the meantime, for CTAR RMSD value rise up to 30 Å indicating structural changes, but the simulation converges to a stable state (Figure 4). The RMSF value of 15 Å also suggest that the CTAR regions experience significant structural flexibility. The Rg values for CTAR varied but were around 20 Å, and one of the triplicate runs (Replicate 2) showed a downward graph trend, suggesting that the protein is becoming more compact or structurally constrained or the simulation has converged (Figure 4c). The CTAR regions experience structural flexibility, possibly indicating functionally significant regions like protein-ligand binding interfaces or enzyme active sites.

**Figure 4.**
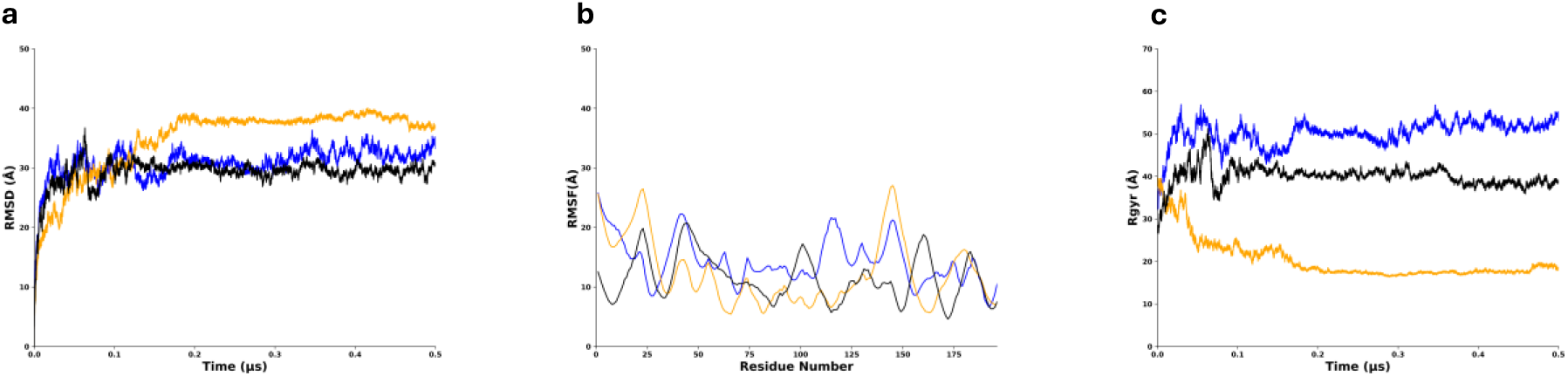
Behaviour of C-Terminal region of LMP1 in MD simulations with replicate 2 (RMSD and Rgyr) has a higher threshold from replicates 1 and 3. Replicate 1 is in blue, replicate 2 is in orange, replicate 3 is in black. **(a)** Root mean square deviation (RMSD) graph. **(b)** Root mean square fluctuation (RMSF) graph. **(c)** Radius of gyration (Rgyr) plot.

Meanwhile, the MD simulation revealed that the RMSD analysis in all the three replicates experience fluctuation between 0.1 µs to 0.3 µs indicating structural deviations and suggesting reduced flexibility in the LMP1 bilayer protein (Figure 5a). Interestingly, the Rg for LMP1 protein became more compact and stable as time passed, ranging from 35–45 Å for all three data. (Figure 5c). RMSF analysis allowed us to identify that the increased flexibility in the LMP1 protein was mainly associated with the CTAR region (Figure 5b). One of the regions involves the residues from Gly300 until residue Asp396, whose average RMSF values were about 30 Å for all three data obtained. The RMSF value confirmed the flexibility structure in the CTAR regions, as stated by previous studies.^28-29^ The studies by Izumi *et al*. and Mainou *et al*. emphasise the complex relationship of CTAR regions in not only B-lymphocyte growth transformation but also its unique role in mediating various signalling pathways such as NF-kB and c-Jun N-terminal kinase (JNK) pathways.

**Figure 5.**
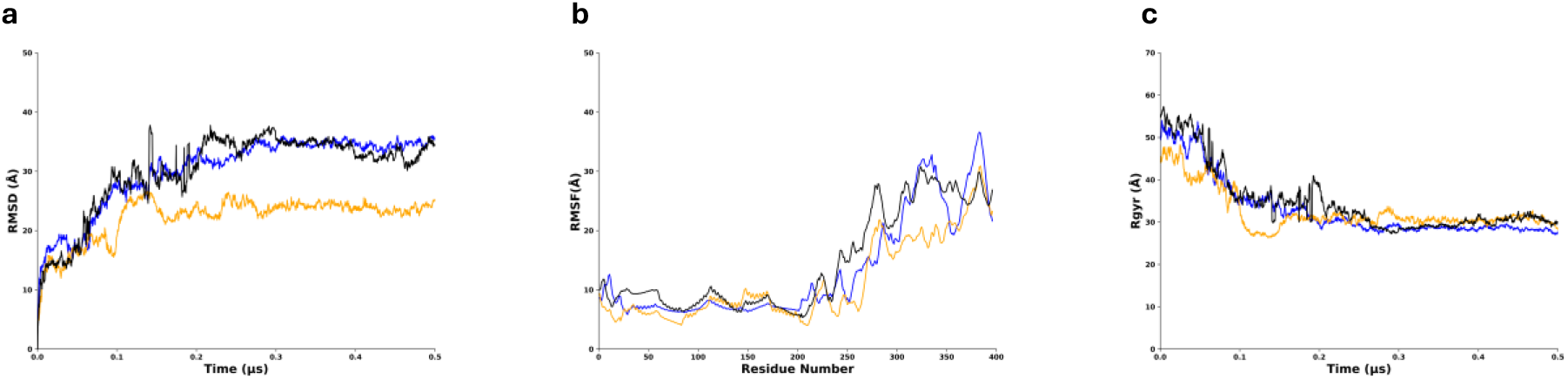
Behaviour of LMP1 bilayer in MD simulations showing replicate 2 (RMSD) has a lower threshold from the other two replicates. Replicate 1 is in blue, replicate 2 is in orange, replicate 3 is in black. **(a)** Root mean square deviation (RMSD) graph. **(b)** Root mean square fluctuation (RMSF) graph. **(c)** Radius of gyration (Rgyr) plot.

The presence of intrinsically disordered regions contributes to the flexibility and adaptability of LMP1, allowing it to interact with diverse binding partners and exert its varied functions. Overall, LMP1 would not be classified as a purely intrinsically disordered protein. It contains structured domains alongside regions exhibiting intrinsic disorder, contributing to its unique functional properties.^15^ Considering that some specific functions of intrinsically disordered regions within LMP1 are still under investigation, the intrinsic disorder of LMP1 can be affected by factors such as post-translational modifications or binding interactions. Recent studies utilising the proximity-dependent biotin identification (BioID) method have revealed a complex interactome associated with LMP1, identifying over 1,200 proteins that interact with LMP1 in various capacities, including direct, transient, or proximal associations.^30^ Among these proteins, several are known to interact with the disordered regions of LMP1, particularly the C-terminal activating regions (CTARs). For instance, TRAF proteins (TRAF1, TRAF2, TRAF3, TRAF5, and TRAF6) are crucial for LMP1 signalling and are known to bind to specific CTARs, facilitating the activation of downstream signalling pathways such as NF-κB and JNK.^31^ The interactions of LMP1 with these TRAF proteins are particularly significant as they help to mediate the oncogenic effects of LMP1 in B-lymphocytes.^32^

### 2.3. Intramolecular Hydrogen Bonding Analysis

Intramolecular hydrogen bonding is important in stabilizing both the secondary and tertiary structure of proteins, contributing significantly to their overall conformational stability and native state.^30^ The hydrogen bonding pattern is a key determinant of the final 3D structure adopted by the polypeptide chain. In this study, the LMP1 predicted structure was simulated for 0.5 µs and we used the cut-off value of more than 60% for all the the simulation runs (Figure 6).

**Figure 6.**
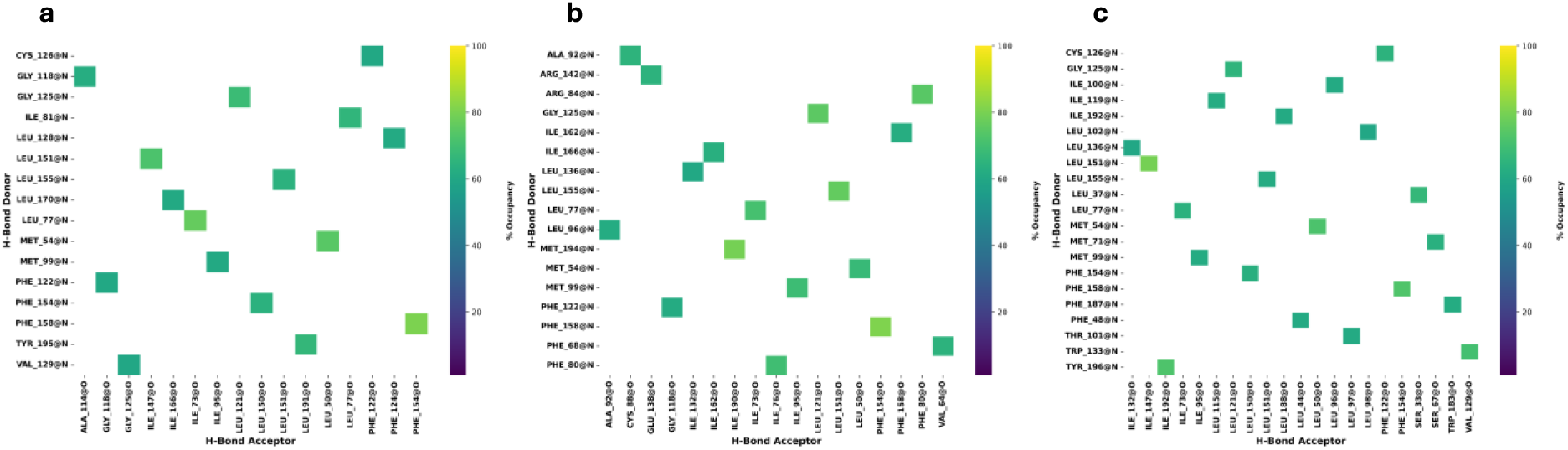
Intramolecular hydrogen bonding analysis for LMP1 Bilayer triplicate data runs with a cut-off limit of 60%. **(a)** Replicate 1 **(b)** Replicate 2 **(c)** Replicate 3.

Our analysis identified some fundamental interactions to explain the minimum ensembles of conformations. Six stable hydrogen bonding interactions namely between Phe154 and Phe158, Ile73 and Leu77, Leu50 and Met54, Leu121 and Gly125, Leu151 and Leu155, and Ile95 with Met99 were found in all the triplicates run. Looking into the residue number position for all of these interactions, we can deduced that all of the stable hydrogen bonding interactions occur in the hydrophobic TMD of LMP1 (between amino acids 25–187). To date there is no direct information available regarding hydrogen bonding in the TMD of LMP1. Hence, further experimental investigation needs to be conducted to provide insights into the presence, strength, and spatial arrangement of TMD intramolecular hydrogen bonding. TMD comprises of approximately 160 amino acid residues that traverse the membrane and contribute to oligomerisation and signal transmission (Supplementary Figure 2a). TMD possesses the innate ability to form homo-oligomers, which may be detected as LMP1 patches in the membrane with the TMD5 playing a particularly important role in the oligomerization and activation of LMP1.^31-32^ Recent research has demonstrated that an intermolecular interaction between TM3-6 and an FWLY motif in TMD1 promotes oligomerisation and NF-kB signaling.^33^ In addition, TMD forms a stable hairpin structure that anchors the protein to the membrane.^34^

### 2.4. Principle Component Analysis and Free Energy Landscape

We used principal components analysis (PCA) and free energy landscape (FEL) to analyse our LMP1 protein further as both analyses offer a comprehensive view of protein structure, dynamics, and stability. PCA identifies the key structural changes, while FEL provides the energetic driving forces behind these changes. The supplementary figure 3 graph displayed the combined data for all of our simulation replicates suggested that there was less variance among all the replicate as the points converged towards the end of the time frame. This indicates that they are all similar in terms of the underlying features used in the PCA. Based on the PCA, the free energy landscape (FEL) plots were projected to identify the preferable conformations of the LMP1 protein (Figure 7a-c). The combined landscape appears to have three visible energy basins. These plots represent relatively stable states of the protein. A ridge in the middle separates the two groups of plots, suggesting a transition state or barrier the protein has to overcome to switch between the two stable states. The blue regions of the free energy indicate a more stable region; meanwhile, the yellow areas depict the less stable areas.

**Figure 7.**
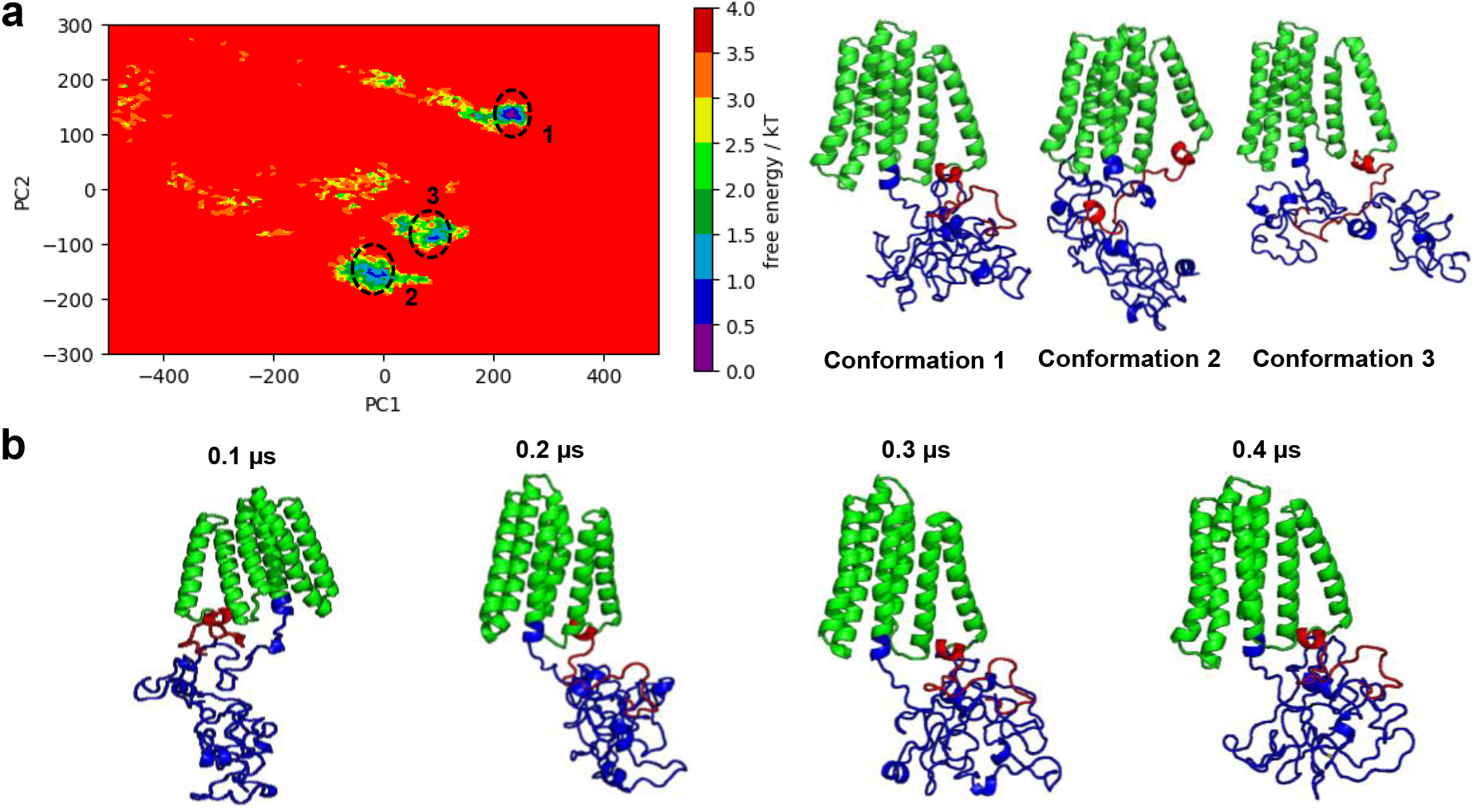
The combined FEL plot with the corresponding conformations of the three basins and the time evolution of LMP1 from replicate 1 that showed the steepest basin. **(a)** The combined FEL plot showed 3 basins which are Conformation 1 (replicate 1), Conformation 2 (replicate 2) and Conformation 3 (replicate 3). **(b)** The time evolution of LMP1 from replicate 1 that showed the steepest free energy basin. The TMD, NTER and CTAR domains are shown in green, red and blue, respectively.

Our PCA and FEL highlighted the analysis obtained from the RMSD and RMSF, whereby the plot showed the stable and less stable regions depicted by the blue and yellow, respectively. The structure of LMP1 is highly dynamic, and it can undergo conformational changes that regulate its activity. Meanwhile, the RMSD and Rg values implied that LMP1 bilayer protein structure has a reduced flexibility and compact structure. We observed that NTER remains stable throughout all the simulation runs meanwhile CTAR region folded nearer to the TMD at the end of the simulation (Figure 7d-f).

Studies have revealed that LMP1 is a highly flexible protein that can adopt different conformations depending on its binding partners and the cellular environment. The stability of NTER of LMP1 (Figure 7d-f) is essential for protein degradation via the ubiquitin-proteasome signalling pathway and cytoskeletal machinery interaction.^35-36^ The NTER of LMP1 affects its half-life and membrane insertion. If deleted, LMP1’s activity and cytoskeleton connection are abolished, and a positive net charge is needed for correct membrane insertion.^37-38^ A potential SH3-binding domain is also found between residues 9 and 20, whereby mutations in this location alter LMP1 patching and reduce EBV’s ability to transform human primary B cells.^39^ However, no protein-binding partners have been identified in LMP1’s NTER domain.

Meanwhile, the conclusion of our MD simulation showed a crumpled CTAR domain structure (Figure 7d-f). CTAR1 domain consists of amino acids 194–232, whereas CTAR2 is placed between amino acids 351–386. The lesser-known CTAR3 domain (amino acids 275-330) is also located between CTAR1 and CTAR2, with few known interaction partners.^40^ The CTAR of LMP1 contains multiple functional motifs involved in signal transduction. These motifs allow LMP1 to activate various signalling pathways, including the NF-κB pathway, the JAK/STAT pathway, and the MAPK pathway.^41-43^ The CTAR domain (amino acids 187–386) attracts TRAFs and TRADD via PQQAT and PVQLSY sites, activating host cell signalling pathways.^44-45^ The chains turn and coil at these PQQAT and PVQLSY sites, as demonstrated by our LMP1 protein molecular simulation (Supplementary Figure 2b). It was also found that CTAR1 shares similarities with CD40’s PxQxT motif, which interacts with TRAF1-3 via the PVQET sequence.^46^ Meanwhile, point mutations in LMP1’s extreme carboxy terminus identified the core motif for CTAR2-mediated NF-kB activation.^47^

## 3. Conclusion

This work continued our previous work and presented an informational structure on EBV LMP1 using accelerated molecular dynamic simulation. We have employed other protein structure websites, such as AlphaFold2 and GalaxyWeb, to confirm our structure better. With little to no known full-length structure of LMP1, our study can be the stepping stone for more research, particularly in the cancer drug discovery field. With convenient and user-friendly web tools such as CHARMM-GUI, we were able to generate the POPC lipid bilayer membrane for the LMP1 for molecular dynamic simulation. The protein simulation produced a free-energy landscape that displayed conformational diversity, particularly in the CTAR region. It revealed that our LMP1 protein structure has a compact structure and reduced flexibility. Despite the diversity, the generated LMP1 conformational ensemble produced states that showed compacted folds as in the TMD region. Our hydrogen bonding analysis also identified some exciting interactions to explain the minimum ensembles of conformations. Although there were some differences in the hydrogen bonding results among our triplicate runs, we have addressed the possible reasons for these discrepancies. The structure of LMP1 is highly dynamic, and it can undergo conformational changes that regulate its activity. Understanding the entire structure of LMP1 is essential for developing new therapies for EBV-associated malignancies. By targeting specific domains or functional motifs of LMP1, it may be possible to create drugs inhibiting its oncogenic activity. This study demonstrates that with further experimental validation of structure-functions relationship, our predicted full-length LMP1 protein structure has the potential to be used for future drug discovery studies.

## 4. Computational Methods

### 4.1. System setting and modelling

The complete LMP1 sequences (386 amino acids) were retrieved from UniProt.^48^ The LMP1 protein used here was designed based on our previous study.^16^

### 4.2. Protein structure prediction website

Some widely used protein structure prediction tools, such as Galaxy Web and AlphaFold2, were used to re-confirm the LMP1 predicted protein structure we designed. GalaxyWeb has a web server specifically designed to predict the structure of protein homo-oligomers called Galaxy Homomer. These are proteins that are formed by the assembly of identical subunits. It uses two approaches, namely template-based modelling and ab initio docking. In our LMP1 predicted structure, the Galaxy Homomer uses the ab initio technique.^49^ Meanwhile, Alphafold2 generated ten PDB files based on credibility ratings to assess the anticipated model’s quality. One was the ‘predicted local distance difference test’ (pLDDT) score, which measures the accuracy of the prediction. The pLDDT is a key metric used by AlphaFold2 to assess the confidence of its protein structure predictions. The pLDDT scores range from 0 to 100, where higher values indicate greater confidence in the accuracy of the predicted residue structure. pLDDT values above 90 suggest incredibly high precision in the predicted structure. Values between 70 and 90 suggest good accuracy or moderate confidence. Meanwhile, values ranging less than 70 reflects poorer accuracy or less reliable predictions.

### 4.3. Lipid membrane modelling

The initial lipid membrane structures were built using the CHARMM-GUI Membrane Builder.^50^ It is a web-based graphical user interface (GUI) for generating input files for the LMP1 lipid bilayer. 1-palmitoyl-2-oleoyl-phosphatidylcholine (POPC) is the lipid bilayer membrane (Supplementary Figure 4). It is a zwitterionic molecule and a neutral phospholipid commonly used in molecular dynamics (MD) simulations to model biological membranes because it is a significant component of cell membranes and forms stable bilayers.^51^ In this work, we did not palmitolylate the LMP1 structure. While palmitoylation is important for LMP1’s localization^52^, and that LMP1 can still associate with lipid rafts even when palmitoylation is disrupted, albeit potentially with altered efficiency or functionality.^53^ This suggests that while palmitoylation enhances LMP1’s ability to interact with lipid rafts, other mechanisms may also contribute to its localization and function. Moreover, POPC lipids are the predominant lipids found in the organelles of mammalian cells, such as the plasma membrane, endoplasmic reticulum, and Golgi apparatus.^54^ This abundance enables a highly accurate representation of the local environment of the peptides within the lipid bilayer, especially when the system is surrounded by an adequate number of water molecules.^55^ The CHARMM-GUI membrane-generated output files in the parm7 and rst7 formats were then used for the protein simulation in AMBER.

### 4.4. MD simulations

One of the first steps necessary to simulate aMD is to do a classical molecular dynamic (cMD) simulation to obtain the average total potential energy threshold (EthreshP) and average dihedral angle energy threshold (EthreshD), as these parameters are needed as input for the aMD simulation. LMP1 EthreshP and EthreshD triplicate run averages were -512,163.333 kcal/mol and 6,428 kcal/mol, respectively. The AMBER 20 MD package performed an adapted LMP1 MD simulation based on AMBER’s accelerated MD tutorial.^56^ For conventional MD simulation, the preparation stages of minimisation, equilibration and heating were done using ff99SB, lipid21, and TIP4PD force field. The Particle mesh Ewald (PME) cutoff distance was kept at 10 Å, and the restraint force was held at 300 kcal/mol. Subsequently, heating was carried out for 20ps, gradually increasing the temperature from 0K to 300K along the NVT ensemble. During the equilibration stage with a time step of 500ps, we employed the NPT ensemble followed by the preparatory production stage for 5ns. Additional parameters were calculated for aMD simulation:-the EthreshP, average total potential energy threshold; alphaP, inverse strength boost factor for the total potential energy; EthreshD, average dihedral energy threshold; and alphaD, inverse strength boost factor for the dihedral energy. The aMD production stage was carried out for 0.5µs and in triplicates (Replicate 1,2 and 3).

### 4.6. Analysis

Analyses were performed using CPPTRAJ on AMBER 20 for trajectory analysis, while VMD (Visual Molecular Dynamics) and PyMOL were used to visualise and check the molecular dynamics simulations in real-time and post-processing.^57-59^ Principal Component Analysis (PCA) was performed by calculating the covariance matrix and diagonalising it to identify the principal components (PCs) and their corresponding eigenvalues. The top PCs represent the most significant variations in protein motion. We estimate the free energy from PCA by projecting the protein coordinates from each trajectory frame onto the top few PCs (usually PROJ1 and PROJ2). This reduces the dimensionality of the data for analysis. Boltzmann distribution is used to estimate the free energy:

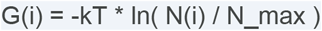

where:

G(i) is the free energy at point i in the PC space. k is Boltzmann’s constant. T is the absolute temperature of the simulation. N(i) is the probability density of finding the protein at point i. N_max is the maximum probability density in the data set.

Free Energy Landscape (FEL) was visualised by constructing a 2D contour plot using axes of the first two principal components (PROJ1 vs PROJ2). Regions with lower free energy values correspond to more populated and stable protein conformations, while higher values indicate less probable and potential transition states.

## Data Availability Statement

All molecular dynamics simulations were conducted using the Assisted Model Building with Energy Refinement (AMBER;version 20.0) using a free academic license. Data are available from the corresponding authors on reasonable request.

### Author Information

Corresponding Authors:

**Taufiq Rahman** − Department of Pharmacology, University of Cambridge, Cambridge CB2 1PD, United Kingdom; orcid.org/0000-0003-3830-5160

**Xavier Chee Wezen** − Faculty of Engineering, Computing and Science, Swinburne University of Technology Sarawak, Kuching 93350, Malaysia; orcid.org/0000-0001-8497-5953

Author

**Dayang-Sharyati D.A. Salam** − Faculty of Engineering, Computing and Science, Swinburne University of Technology Sarawak, Kuching 93350, Malaysia; orcid.org/0000-0002-6201-4227

## Author Contributions

DSDAS: conception and design of the work; analysis and interpretation of data; drafted the work. KKJG: interpretation of data; revised and editing. HSS: revised and editing. IRHG: revised and editing. TR: revised and editing. XCW: interpretation of data; revised and editing.

