## Supplementary Information for "*In silico* modelling and characterization of Epstein–Barr virus LMP1 protein"

### *Supplementary Table and Figures*

**Supplementary Table 1 Ab initio docking results for LMP1 predicted structure.**

| Model No. | No. of Subunits | Interface Area (in Angstrom $^2$) | Docking Score |
|---|---|---|---|
| 1 | 2-mer | 2279.7 | 2350.543 |
| 2 | 2-mer | 2173.8 | 2206.530 |
| 3 | 2-mer | 1804.2 | 2103.901 |
| 4 | 5-mer | 7891.9 | 1012.986 |
| 5 | 3-mer | 3869.4 | 914.557 |

a

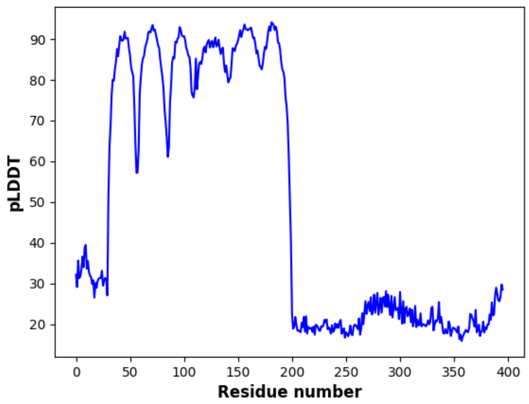


b

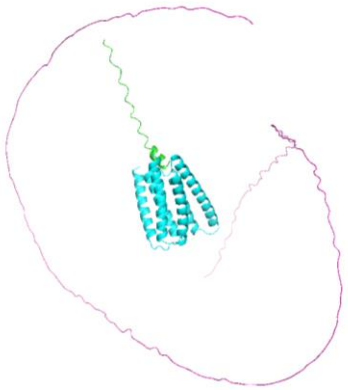

**Supplementary Figure 1 (a)** Per-residue confidence score for LMP1 model 3 **(b)** AlphaFold2 LMP1 model 3 image

a

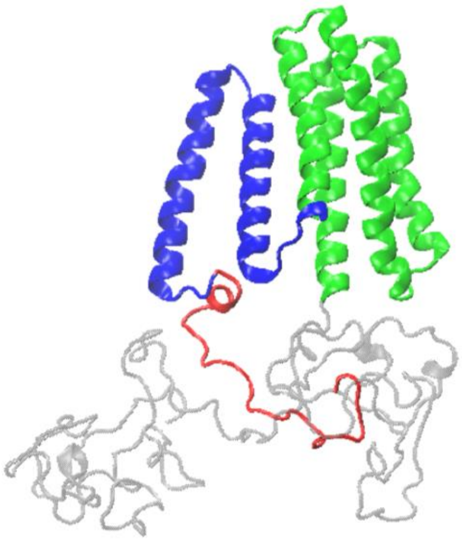

b

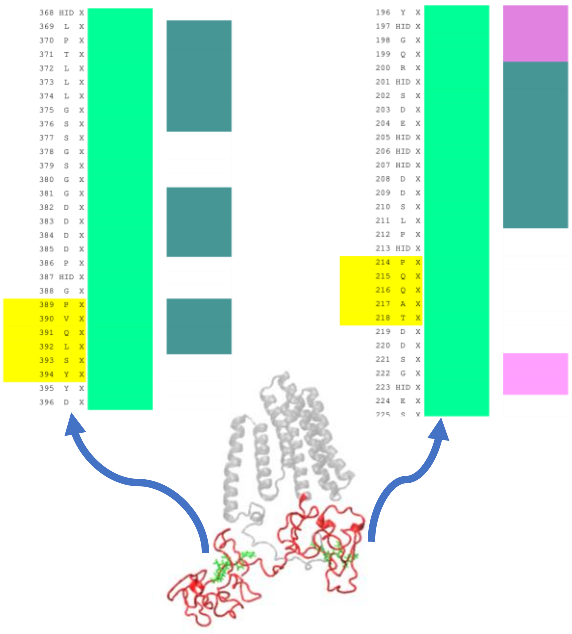


**Supplementary Figure 2 (a)** The molecular image of TMD3-6 (green colour) and TMD1-2 (blue colour) responsible for the oligomerisation of the LMP1 protein. The red-colour chain represents the N-terminal domain of the LMP1 protein. **(b)** The CTAR1 and CTAR2 domains activate the cells.

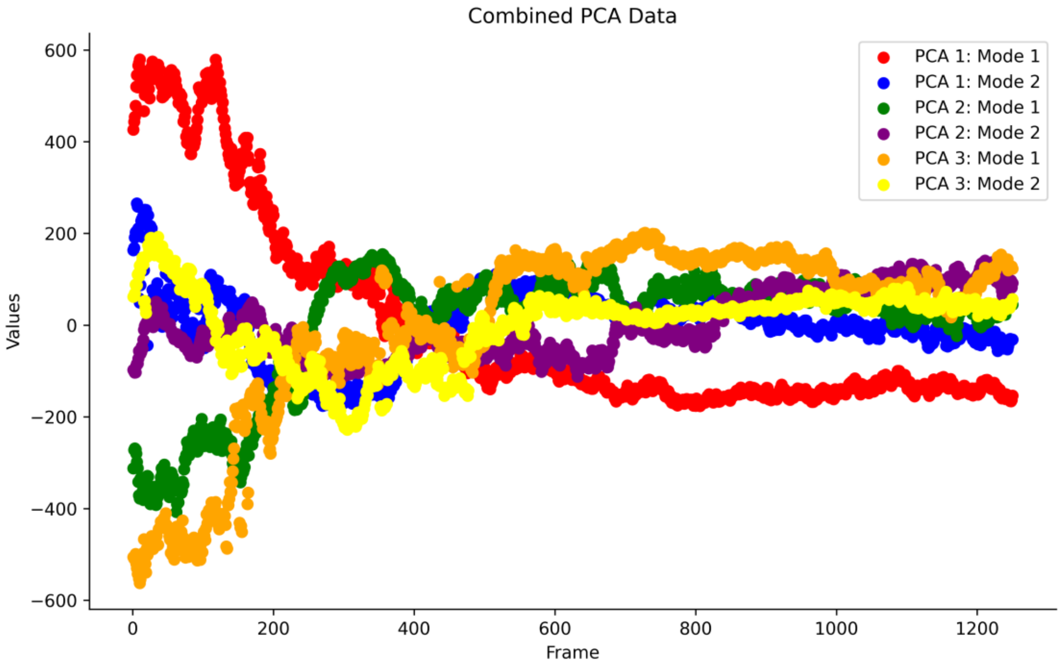


**Supplementary Figure 3** Graph shows combined PCA data for all the triplicate runs with the values on the y-axis and frame on the x-axis.

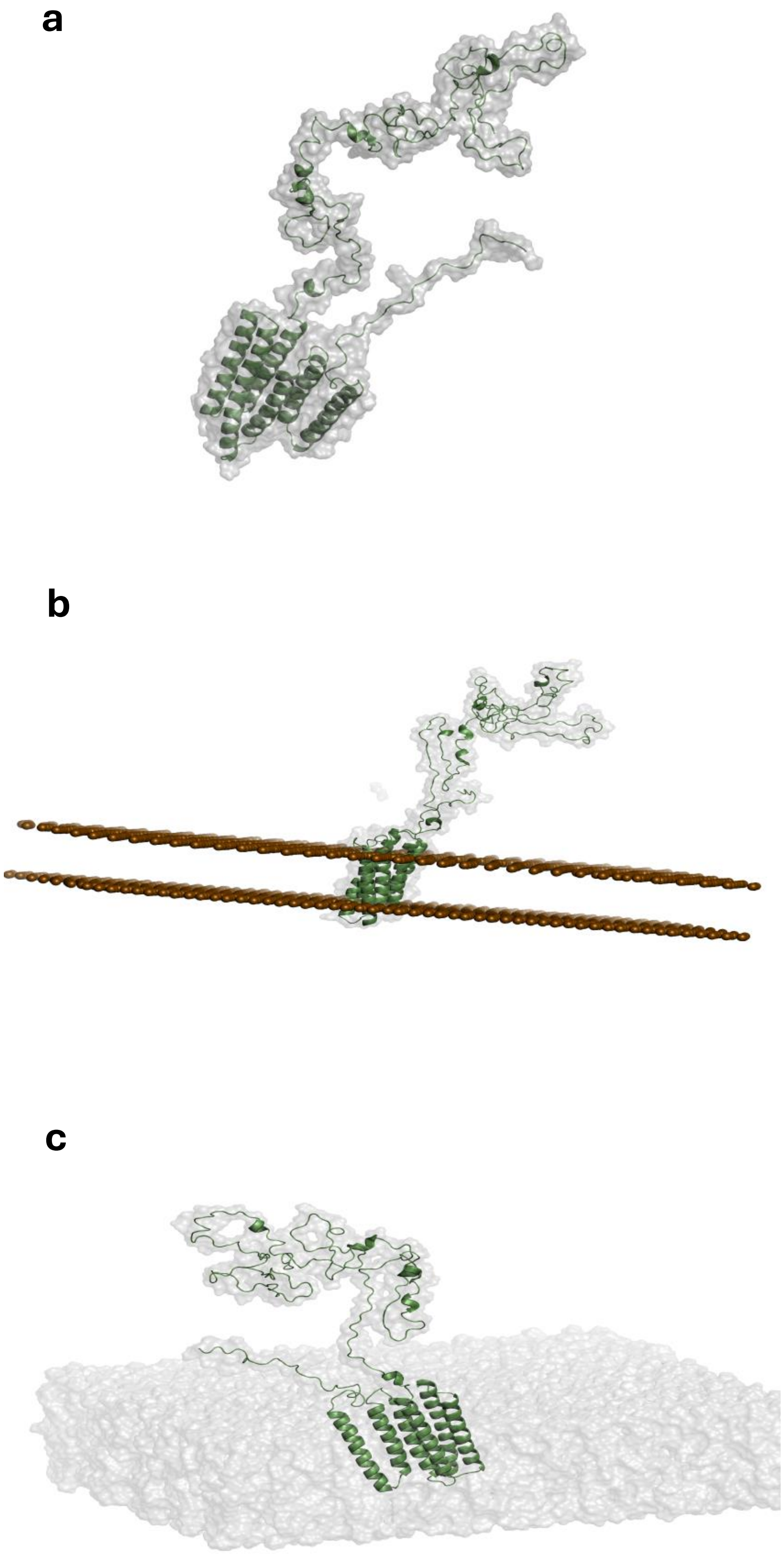
a
b
c

**Supplementary Figure 4** Molecular graphics of LMP1 membrane protein structure built with POPC lipid bilayer **(a)** STEP 1 **(b)** STEP 3, and **(c)** STEP 5 in CHARMM-GUI Membrane Builder. Water molecules and ions are not shown in (c).
